# pH Induced Changes in Protein Structure and Hydration

**DOI:** 10.64898/2026.05.13.724817

**Authors:** Anusree Sen, Rajib Kumar Mitra, Jaydeb Chakrabarti

## Abstract

The molten globule (MG) state is an intermediate in the unfolding pathway of proteins, typically triggered by denaturing agents such as urea, extreme pH, high pressure, or heat. The microscopic details of such states are far from understood. Here we study the MG states in protein Hen Egg-White Lysozyme (PDB ID: 1AKI) using microscopic constant pH molecular dynamics (CpHMD) simulations and experiments across a wide pH range. We observe that the titratable residues act as key drivers of conformational fluctuations, promoting the emergence of MG states at extreme pH. These states display partial unfolding, and small global structural changes (*<* 7% deviation). Hydration around the fluctuating acidic residues shows reduced water density and weakened hydrogen bonding at low pH. At high pH, hydration around acidic residues increases relative to pH = 7, whereas hydration around basic residues decreases. The translational and rotational dynamics of hydration water also exhibit pronounced pH dependence: the translational diffusion coefficient (*D*_trans_) increases linearly with decrease in pH in acidic medium and increases linearly with increasing pH in the basic regime. The rotational diffusion (*D*_rot_) shows similar dependencies on pH except a break at pH ≈ 4 corresponding to acidic residue p*K*_*a*_ values. Our results may be useful to identify ligand binding of lysozyme in extreme pH conditions.

## 1. Introduction

Many proteins exhibit partial unfolding, while retaining their overall tertiary structure in near denaturing conditions,^1^ achieved by elevating temperature, addition of denaturing agents like Urea, changing pH and so on.^1–3^ Experimentally the partial unfolding has been observed for proteins, like *α*-lactalbumin,^4^ IgG, papain, SP-C and Ubiquitin^5^ to name a few. Computer simulations show only local structural changes while the overall global structure in a partially unfolded state of a protein remain largely intact. This is also known as a molten globule(MG) state^6^ of the protein. The nature of structural fluctuations and the hydration in MG states remain poorly understood.

pH of a medium modifies the charged states of titrable residues in a protein. This results in changes in the electrostatic interactions in the system, and consequently, affects the protein structure and hydration substantially. Hen Egg-White Lysozyme(HEWL) ^7^ is a stable antibacterial protein, consisting of 129 amino acids and has a compact regular structure. The secondary structural segments of the protein consists of four *α*-helices, three 3_10_-helices, and three standard *β*-sheets.^8–10^ This protein is linked to systemic non-neuropathic amyloidosis diseases where large amount of lysozyme aggregates under extreme pH conditions, leading to organ failure.^11,12^ Numerous experimental studies on lysozyme using various spectroscopic techniques including THz, CD, and 2D-IR under high temperature and pH conditions away from the neutral condition^13–15^ establish partial unfolding and changes in the secondary structural elements. An earlier study using the Hydrogen exchange measurements shows that the partially unfolded state of lysozyme at pH = 2 retains a compact hydrophobic core with native like interactions. ^16^ This reduces the ability of the protein to break down bacterial cell walls, thereby affecting its antibacterial activity.^7^ Hen egg-white lysozyme (1AKI) exhibits maximum anti-microbial and enzymatic activities in the pH range 6–9, enhancing ligand binding and enzymatic activity.^17^ Under acidic pH and thermal stress (e.g., pH = 2 and 65°C), lysozyme adopts a *β*-sheet rich partially unfolded conformation, which promotes aggregation.^18^ Experiments also show that at alkaline pH and in presence of tertiary butanol, lysozyme adopts a partially unfolded state.^19^ Atomistic molecular dynamics(MD) simulations report faster partial tertiary unfolding with retained secondary structure and slightly increased *β*-sheet content at acidic pH.^20,21^ 2D-IR spectroscopy combined with MD simulations^22^ show increased solvent accessible surface area and partial structural destabilization under acidic conditions. A recent study using Raman spectroscopy and MD simulations using molecular docking show that disulfide bond breakage at pH = 12.2 leads to *α*-helix unfolding, *β*-sheet formation, and stable aggregate formation through intermolecular interactions.^23^ Many simulation based investigations including unfolding trajectories, long time MD, and enhanced sampling studies have established the idea that lysozyme can populate compact, partially unfolded intermediates reminiscent of MG ensembles under denaturing conditions.^24,25^

The conventional MD simulations assume fixed protonation states and cannot fully capture pH-dependent behavior at the microscopic level. ^26^ CpHMD^27^ simulations with explicit solvent allows dynamic titration of ionizable residues. CpHMD is a hybrid simulation scheme where, MD propagation of atomic coordinates is coupled with Monte Carlo(MC) sampling of the protonation states of titratable residues in order to maintain the system at a specified pH. CpHMD simulations have been applied to lysozyme earlier to compute the titration curves.^28^ This technique has been applied to understand MG state in milk protein, *α*-lactalbumin which forms MG states in acidic medium and acts as carrier of charged fatty acids.^29^ The simulations show that the protein undergoes conformational changes in localized regions which help the binding of fatty acids. While extreme pH can destabilize lysozyme into a partially unfolded structure it is unclear whether such conditions induce a MG state with only localized structural changes.

Hydration of proteins refers to the interaction between protein molecules and surrounding water molecules. Water forms a dynamic layer around the protein surface, stabilizing its structure through hydrogen bonding and electrostatic interactions. This hydration shell plays a crucial role in maintaining protein folding, flexibility, and biological function. Although the solvation properties of small molecules^30,31^ and structured proteins^32,33^ are well understood, gap exists in understanding how pH dependent protonation leads to reorganizes the hydration environment of lysozymes under extreme pH. We perform microscopic simulations on HEWL (PDB id: 1AKI)^34^ using the CpHMD simulations across the entire pH range (1–13). Our primary focus is specifically on structure and hydration surrounding the titrable residues near denaturing conditions at extreme pH.

The key results in our studies are as follows: (1) The protein gets extended at extreme pH. While the overall secondary structure of lysozyme remains nearly intact, local changes occur particularly around the titrable residues with varying pH as seen in experiments, machine learning analysis and conformational thermodynamics data. These observations confirm a MG state of the protein. (2) The fluctuating acidic residues show lower water density and fewer hydrogen bonds at low pH. In contrast, at high pH, hydration around the acidic residues further increases compared to pH = 7, while hydration around the basic residues decreases significantly. (3) We find that the dynamics of hydration water are strongly modulated by pH. The translational diffusion of water increases as pH decreases under acidic conditions and increases with pH in basic condition. The rotational diffusion follows a similar trend but exhibits a distinct change in behavior around pH ≈ 4. These variations are closely linked to pH induced fluctuations in the protein and the surrounding hydration shell. Some of the structural results corroborate well with our zeta potential and mid-IR measurements. The paper is organized as follows: The system details, experimental and simulation details and the analysis given in Section 2. Section 3 gives the simulation and experimental results. The paper is concluded in section 4.

## 2. Methods

### 2.1 Simulation details

We perform CpHMD simulations following the hybrid MD – MC scheme described by Swails et al.^35^ using the AMBER simulation package. During protonation state change attempts, the electrostatic component of the potential energy is evaluated using the Generalized Born (GB) implicit solvent model.^36^ Protonation states of titratable residues are randomly modified, and the resulting change in electrostatic free energy is used in a Metropolis criterion to determine whether the new protonation state is accepted or rejected. If a protonation state change is accepted, the solute coordinates are temporarily held fixed while MD is performed on the solvent to allow relaxation around the updated protonation state. During the MD propagation itself, however, the system is simulated in explicit solvent.

In this work, CpHMD simulations are performed at discrete pH values ranging from pH = 2 to pH = 13. Specifically, simulations are carried out at pH = 2, 3, 4, 5, 6, 7, 8, 9, 10, 11, 11.5, 12, 12.5, and 13. The AMBER package defines titratable side chains for the residues aspartate (ASP), glutamate (GLU), histidine (HIS), lysine (LYS), arginine (ARG), tyrosine (TYR), and cysteine (CYS).^37^ This protein contains 4 disulfide bonds formed by eight CYS residues, resulting in four cystine pairs. To correctly represent these disulfide linkages during system preparation, all cysteine residues involved in disulfide bonding are defined as CYX in the topology generation step. TYR has not been titrated because its protonation/deprotonation is slow to sample and highly environment-dependent, making simulations less stable and harder to converge. Under acidic conditions (pH = 2 – 6), the acidic residues ASP and GLU are titrated because their p*K*_*a*_ values are approximately 4.0 and 4.3, respectively.^38^ Near neutral pH (pH = 7), histidine (HIS), with a p*K*_*a*_ value of approximately 6.5, participates in the titration process and gets deprotonated above this point till the maximum pH we study(pH = 13). Under basic conditions, basic residues such as lysine (LYS) and arginine (ARG), which have p*K*_*a*_ values of approximately 10.5 and 12.5 respectively,^39^ are allowed to titrate. Additional simulations at pH = 11.5 and 12.5 are performed to better capture the behavior of the system under strongly basic conditions. Since CpHMD in AMBER does not currently support titration of Nor C-terminal residues, these termini are excluded from the titration.

The simulations are carried out using the AMBER software in an orthorhombic simulation box of size 65.71 Å × 70.41 Å × 61.63 Å. We use the crystal structure of lysozyme (PDB ID: 1AKI) as the starting configuration.^34^ The total number of atoms inside the box is 22459. 20454 water atoms are added in the box. 8 Cl^−^ ions are randomly added to neutralize the total charge of the system at pH = 7. The number of particles depends on the overall charge state of the protein where ions are added to maintain the electrostatic neutrality at a given pH. At each pH, the protonation states of ASP, GLU, ARG, LYS, and HIS are assigned using p*K*_*a*_ values reported in the AMBER CpHMD tutorial.^40^ Protein interactions are modeled using the AMBER FF10 force field, ^41^ and the TIP3P water model is employed.^42^ A 10 Å cutoff is used for van der Waals interactions, and the Particle Mesh Ewald method^43^ is used for electrostatics. The energy is minimized using 5000 steps of steepest descent and conjugate gradient methods with a 10 kcal·mol^−1^·Å^−2^ positional restraint on the backbone atoms. This is followed by a heating phase at constant volume from 10 K to 300 K over 1000 ps. The system is equilibrated in the NVT ensemble for 30 ns at 300 K using the Nosé–Hoover thermostat. ^44^ Periodic boundary conditions are applied in all three directions. We perform 1 *µ*s simulations. Coordinates are saved every 2 ps. A multiple time step algorithm is used to integrate the equations of motion, and bonds involving hydrogen atoms are constrained using the SHAKE algorithm.^45^ We take three independent runs for each pH. For analysis, the equilibrated portion of the trajectory is divided into *N*_ind_ = 50 windows. For each quantity of interest we compute the mean over each window, and the overall average is taken as the mean of the window means. The error in the mean is given by

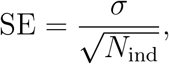

where *σ* is the standard deviation of the distribution of the window means.

After the CpHMD simulations reach equilibrium at each pH, representative configurations with the corresponding equilibrium protonation states of the titratable residues are extracted. Additional 4 ns production simulations are then performed with fixed protonation states while saving coordinates every 0.05 ps in order to capture the short-time dynamics of hydration water. This procedure avoids artifacts associated with solvent relaxation steps that accompany protonation-state change attempts during CpHMD simulations.

#### RMSD of the system

The root-mean-square deviation (RMSD) of a protein in solvent is defined as

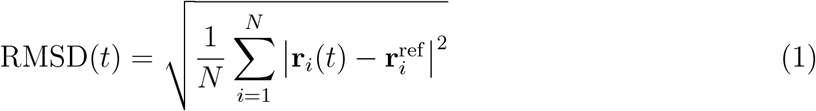

where *N* is the number of selected atoms (typically backbone atoms), **r**_*i*_(*t*) is the position of atom *i* at time *t*, and 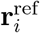 is the reference structure (after optimal alignment to remove rotational and translational motions). We compute the following quantities from our simulated trajectories.

#### Energy Auto-correlation function

The energy auto-correlation function (EACF) of the system is defined as

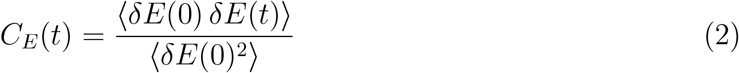

where *δE*(*t*) = *E*(*t*) − ⟨*E*⟩, with *E*(*t*) being the instantaneous energy and ⟨*E*⟩ its time average.

#### Total Charge of the protein

The pH, pK_a_ and *f* (fraction of deprotonation) of a titrable moiety are related by the Henderson-Hasselbalch equation^46,47^

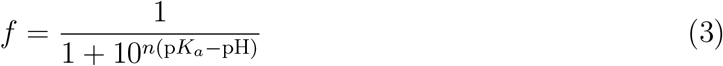

where the *f* is the fraction of de-protonation. *n* is the Hill-coefficient. To get the fraction of protonation we use 1 − *f*.^48^

#### Secondary Structure

The secondary structure analysis is done using **secstruct under cpptraj module** of AMBER^49^ for different equilibrated trajectories at different pH conditions.

#### Radius of Gyration of Protein

The radius of gyration(*R*_*g*_)^50^ has been calculated using the formula shown below:

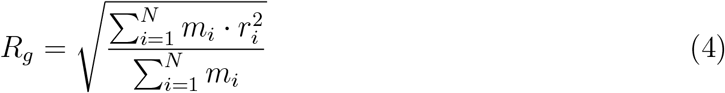

where the sum is over all particles(N). The term *m*_*i*_ represents the the mass associated with the *i*^*th*^ particle in the system. This is the sum of the products of each element’s mass and the square of its distance from the center of mass.

#### Conformational Thermodynamics Analysis

The free energy (Δ*G*) and entropy (*T* Δ*S*) associated with conformational changes in proteins are derived using histograms of dihedral angles. ^51^ The total conformational free energy and entropy costs for each residue can be estimated by summing the individual contributions of free energy and entropy for each dihedral angle within that residue.^52^ The equilibrium conformational free energy change (Δ*G*) between two distinct conformational states, A and B, for a given dihedral angle, *θ*, is calculated as:^29^

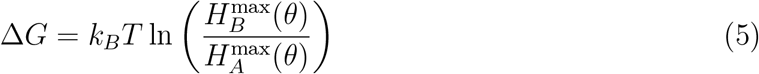

where *H*_*X*_(*θ*) represents the normalized probability distribution for the dihedral angle *θ* in state *X*, and the superscript “max” refers to the peak probability value. These histogram peaks are taken as the equilibrium points for each dihedral angle. To calculate the conformational entropy (*T* Δ*S*) for a particular dihedral angle, the Gibbs entropy formula is used:^53^

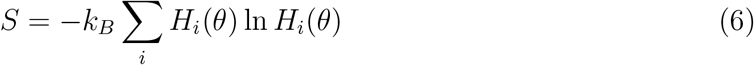

where the sum is taken over all histogram bins *i*. The entropy change (Δ*S*) between states A and B for any dihedral angle *θ* is given by:

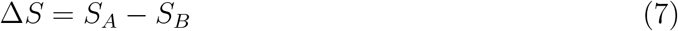

These calculations are performed using in-house software (https://github.com/snbsoftmatter/confthermo). Residues with positive values for both Δ*G* and *T* Δ*S* are labeled as destabilized and disordered. To estimate the error, the equilibrated trajectory is divided into five equal segments, calculating Δ*G* and *T* Δ*S* for each segment.^54–57^ A detailed description of the histogram-based method (HBM) regarding the conformational thermodynamics calculation is reported in literature.^58–60^

#### Dihedral principal component analysis

The protein structure can be effectively characterized by its backbone dihedral angles, *ϕ* and *ψ*. We apply principal component analysis (PCA) using the advanced dPCA+ method, as developed by Sittel et al.^61^ Traditional PCA can occasionally encounter issues when computing the covariance matrix and when projecting circular coordinates onto the eigenvectors. The dPCA+ method addresses these challenges by periodically adjusting dihedral angles to align the largest sampling gap at the periodic boundary, reducing errors associated with the circular nature of dihedrals. This method leverages the fact that, due to steric constraints, dihedral angles do not span the entire theoretical range [−*π, π*] but are limited to certain regions. By applying dPCA+ to molecular dynamics data of a protein, it is possible to obtain the free energy landscape. Based on the structure of a one-dimensional projection of this landscape, additional principal components may be selected for further analysis. In this study, terminal residues are not included in the analysis. We do the dimentional reduction using the dPCA+ method on our system for three different solvent conditions i.e pH = 3, pH = 7, pH = 13.

#### Density based clustering

We perform a robust density-based geometrical cluster analysis^62^ over a landscape in the hyperspace spanned by the dihedral principal components (PCs). For each structure in the trajectory, we count the number of frames within a fixed radius *R* from the given frame inside the hypersphere. Normalizing this count gives the density of sampling probability *P*. The free energy is estimated using the equation:

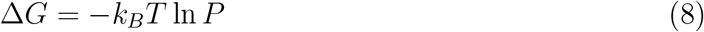

Initially, an energy cut-off value at a relatively low free energy (Δ*G*_0_ ≤ 0.1*k*_*B*_*T*) is defined. All structures below this cut-off are considered, while others are ignored. Frames that are closer than a specified lumping radius (Δ_lump_) are assigned to the same cluster. The energy cut-off is gradually increased in steps of 0.1*k*_*B*_*T* until all clusters converge at the energy barrier. This procedure ensures that all structures obtain a specific cluster membership. According to Nagel et al.,^63^ in most cases, Δ_lump_ itself is a good choice for the clustering radius *R*. Here, we set both *R* and Δ_lump_ equal to 0.52. At the end of the density-based clustering, when all data points are considered, the minimal population *P*_min_ of each state is determined as the percentage of all data points(typically taken between 12 - 15). This prevents the inclusion of small microstates within the same minimum, which could arise due to local free energy fluctuations. Since *P*_min_ affects the number of microstates, it should be chosen based on the desired level of coarse graining.

#### Dynamical clustering

Density-based geometrical clustering is expected to provide a valid primary description of the system when the conformational states of different geometrical configurations are separated by large barriers.^63^ However, sometimes small structural changes are observed. Even in the case of rare transitions between states, geometrically distinct but dynamically close states may be artificially separated. Conversely, in cases of low sampling, dynamically distinct states might be considered geometrically close and, therefore, wrongly grouped together. This error can be minimized by considering a dynamic clustering method that combines MD frames based on their temporal evolution rather than geometrical changes. Here, we employ the most probable path (MPP) algorithm developed by Jain et al.^64^ for dynamic clustering. Initially, given a set of microstates, the transition matrix between these states is calculated. For each state, if the self-transition probability is lower than a certain metastability criterion *Q*_min_ ∈ (0, 1], the state will be merged with the state that has the highest transition probability and the lowest free energy. This process is repeated until no further transitions occur for a given *Q*_min_. At pH = 7, microstate analysis with a threshold *P*_min_ = 15 identified 30 microstates, which are reduced to 14 metastable states using a lag time of *t* = 20 ps, *Q*_min_ = 0.077, and a coring procedure to eliminate spurious transitions. Next, we perform the microstate analysis using a threshold of *P*_min_ = 15, identifying 28 microstates for pH = 3. These were reduced to 12 metastable conformational states by applying a lag time of *t* = 20 ps and a metastability criterion of *Q*_min_ = 0.090, while eliminating spurious transitions via a coring procedure. We analyze dihedral angle fluctuations for pH = 13 using the same methodology as in acidic conditions. The lag time remains *t* = 20 ps, while the metastability criteria, *Q*_min_, is adjusted to 0.085.

#### Essential coordinates (ECs)

We have employed supervised machine learning techniques to identify essential coordinates for meta-stability in the MG state for solvent pH = 3 and pH = 13. Specifically, we use the extreme gradient boosting (XGBoost) algorithm^65^ to identify an ECs of the system in the meta-stable state. In this algorithm, given a trajectory of MD coordinates in terms of the dihedral angles and a meta-stable state obtained from clustering, a machine learning model is constructed by minimizing a loss function. The overall accuracy of the model is evaluated by dividing the available data into training and test sets. The importance of a coordinate is determined by the gain in the loss function value. Any MD coordinate with a higher gain in the loss function is considered more important for characterizing the state than others. Once the model is trained, all the dihedrals are ranked based on their importance. When a non-essential dihedral is discarded, the accuracy of the model does not change significantly. This process allows for the identification of ECs that are critical for discriminating between the states. For this study, all XGBoost parameters are selected as in earlier literature.^66^

#### Density profile calculation

We calculate the density profile of water molecules around the protein, as well as around specific groups of residues like acidic, basic, and hydrophobic under different pH conditions using the following expression:^67^

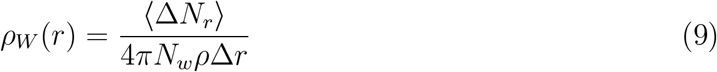

Here, *r* is the distance between the center of mass (COM) of the protein or a selected group of residues (e.g., acidic, basic, or hydrophobic) and the oxygen atom of a water molecule. ⟨Δ*N*_*r*_⟩ represents the average number of water oxygen atoms within the spherical shell of thickness Δ*r*, centered at *r. N*_*w*_ is the total number of water molecules in the system, and *ρ* is the density of bulk water.

Alternatively, the density profile can be computed using a delta function representation:

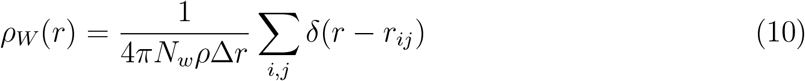

where *r*_*ij*_ is the distance between the COM of the *i*^th^ of the entire protein or a specific residue groups and the oxygen atom of the *j*^th^ water molecule. A hydration site is identified as a local maximum in the resulting solvent density profile near the protein or residue surface.

#### Hydrogen bonds analysis

Geometric criteria for determining these inter protein-water hydrogen bonds is taken from the literature.^68^ The hydrogen bonds are calculated using the MD Analysis Hydrogen Bonds library provided in MDAnalysis Hydrogen Bonds Documentation. ^69,70^ The average number of H-bonds is calculated as:

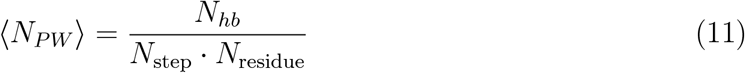

where *N*_*hb*_ is the total number of H-bonds found during the analyzed period, *N*_step_ is the number of trajectory frames, and *N*_residue_ is the number of residues of each type. Here using these criteria we calculate the mean number of hydrogen bonds formed per residue between protein and water molecules surrounding it for different solvent pH conditions.

#### Hydration dynamics

Survival Probability (*S*(*t*)) gives the probability for a group of particles to remain in a certain region. The *S*(*t*) is given by:

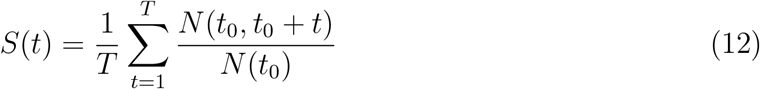

where T is the maximum time of simulation, *N* (*t*_0_) the number of particles at time *t*_0_, and *N* (*t*_0_, *t*_0_ + *t*) is the number of particles at every frame from *t*_0_ to (*t*_0_ + *t*) within the first hydration shell.^71^ The residence time

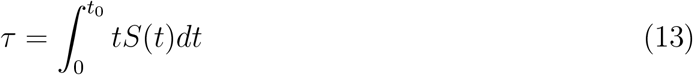

For carrying out the integration we use fitted curve to *S*(*t*).

Next we measure the translational mean-squared displacement (MSD) of hydration water as follows:^72^

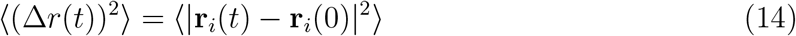

Here, **r**_*i*_(*t*) is the position of water particle *i* at time *t*, and ⟨…⟩ denotes an ensemble average. Calculations are restricted to the first hydration layer (up to 5 Å), with residence times used as a cutoff for MSD calculations. At short times (*τ* < 10 ps), MSDs exhibit a subdiffusive regime, transitioning to linear MSDs in the diffusive regime. The diffusion coefficient *D* is defined in long time domain where MSDs are linear.

Now we consider the water molecules as dipoles, and therefore their electrostatic interaction with the protein is influenced by the charge of the protein hence the solvent pH. The orientation of water molecules at an interface is characterized by the angle *θ* between the dipole vector *µ* (defined as the vector from the water oxygen to the midpoint between the two hydrogen atoms) and the *z*-axis. The rotational dynamics of the hydration water dipoles under different pH conditions can be analyzed using the mean squared angular displacement(MSDs), defined as

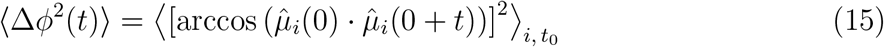

where 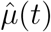 is the unit vector along the dipole axis at time *t*, and the brackets ⟨·⟩ denote averaging over all water molecules and time origins *t*_0_. For rotational diffusion in three dimensions, the MSD grows linearly with time in the long-time diffusive regime:

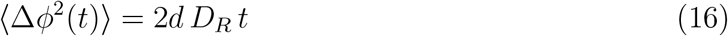

where *d* = 3 is the rotational dimensionality, and *D*_*R*_ is the rotational diffusion coefficient.

### 2.2 Experiments

Lysozyme samples are initially prepared at a concentration of 10 mg/mL in 20 mM His/His–HCl buffer at pH = 6.0. To achieve the desired working concentration, the protein solutions are diluted with ultrapure water to approximately 0.7 mg/mL. Subsequently, the samples are mixed 1:1 (v/v) with the respective sample buffers to yield final solutions suitable for the experimental conditions.

The following buffers are employed to ensure the appropriate ionic and pH conditions for analysis: Acetic acid–sodium acetate buffer: 4.6M molar concentration. KCl–NaOH buffer: 12.6M molar concentration. HCl–KCl buffer: 2M molar concentration. The final protein concentration after dilution and mixing was approximately 0.35 mg/mL, ensuring optimal compatibility with the experimental setup. All buffers are freshly prepared, filtered, and adjusted to the required pH to minimize contamination and ensure consistency. Zeta-potential measurements are conducted using a Nano S Malvern instrument.^73^ The experiments are based on the principles of the Poisson-Boltzmann theory, accounting for the electrostatic double layer surrounding charged particles in an electrolyte solution.^74^ Measurements are performed with the default Henry function settings. Corrections are applied to obtain the zeta potential, which is further rescaled to calculate the surface charge density of the protein.

#### Total charge of the protein for different pH conditions

In Zeta-potential measurements, the surface charge density *σ* of a protein is calculated using the following formula:

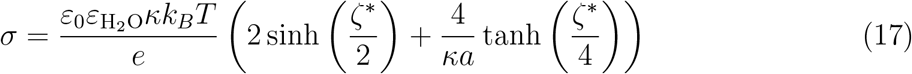

where *ε*_0_ is the vacuum dielectric permittivity, 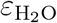 is the dielectric permittivity of water, and *κ* is the Debye-Hückel parameter,^75^ which accounts for the ion screening effect in solution. *k*_*B*_ is the Boltzmann constant, *T* is the absolute temperature, and *e* is the elementary charge. *ζ*^∗^ is the rescaled zeta potential, corrected by the Henry function, and *a* is the radius of the particle.

In the Zeta-potential measurement method, the zeta potential (*ζ*) of a particle in an electrolyte solution is first measured.^76^ This potential reflects the electrostatic interactions between the particle’s surface and the surrounding ions. The Debye-Hückel parameter *κ*, which describes the screening effect of the ions in the solution, is then calculated. To account for the measured zeta potential, a correction is applied using the Henry function, which takes into account the size of the particle and the ionic strength of the solution. The corrected zeta potential *ζ*^∗^ is then used to calculate the surface charge density *σ* of the particle. The equation above incorporates the temperature, the dielectric properties of the medium, the particle size, and the ionic strength of the solution to determine the surface charge density, which is essential for analyzing the behavior of charged particles in the medium. ^77^

#### Overall structural changes of the protein for different pH conditions

In experiments, FTIR measurements are performed in the spectral window of 1000–1800 cm^−1^ for mid-IR regions, using a Vertex 70V (Bruker, Germany) Fourier transform infrared spectrometer with a DLaTGS detector.^78^ The spectrometer is equipped with an attenuated total reflection (ATR) sample compartment featuring a diamond crystal with a refractive index of 2.38. All measurements were conducted in ATR mode.^79^ The sample compartment is thoroughly cleaned and evacuated using a vacuum pump each time prior to measurements. Each measurement is an average of 64 scans with a resolution of 4 cm^−1^. The spectral output, obtained in ATR units, is then converted into absorbance using the following relation:^80^

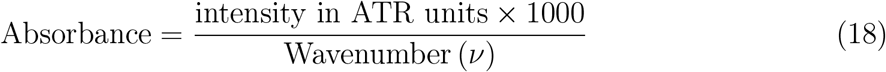

## 3. Results and Discussions

We judge equilibration from the RMSD of the system with respect to the initial frame, as shown in Fig.S1(a). It is observed that, under all pH conditions, the system reaches equilibration after approximately 50 ns. We calculate different quantities of interest by dividing the equilibrated trajectory into independent windows. We compute the energy auto-correlation function of the equilibrated trajectories using Eq.2. The corresponding energy autocorrelation functions for pH = 3, 7, and 13 are shown in Fig.S1(b)-S1(d). The decay is well described by a single-exponential form 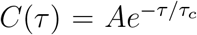. We find *τ*_*c*_ ≈ 0.7 ns. Accordingly, the equilibrated trajectory is divided into 50 blocks each consisting of 10,000 frames each spanning time ≈ 20ns order of magnitude larger than *τ*_*c*_. We compare the estimated pKa values of the titratable residues from the long equilibrated simulation trajectories with the reported experimental measurements^37^ in Table.S1. Overall, the calculated values show reasonable agreement with experiment, demonstrating that the CpHMD simulations adequately reproduce the protonation behavior of the residues for different pH conditions.

All structural and thermodynamic properties reported in the result section (e.g., radius of gyration, secondary structural changes, identification of the ECs, conformational thermodynamics, density profiles and h-bond analysis) are computed from these equilibrated CpHMD trajectories. In contrast, dynamical properties of hydration water are obtained from separate short production simulations with fixed protonation states extracted from the equilibrated CpHMD ensembles, as described in the Methods section.

### 3.1 Changes in protein charges

The titration curves for all titratable residues are shown in Fig.S2(a)–S2(q). The predicted p*K*_*a*_ values, the computed Hill coefficients (*n*), and the experimental p*K*_*a*_ values reported in the literature^37,39,81,82^ are summarized in Table S1. By fitting the titration, *f* vs pH curve obtained from simulations, pK_a_ and n are estimated using Eq.3. The fraction of protonation is given by 1 − *f*. We compute the charge of each residue(Q) at a given pH using *f* as follows. For acidic residues such as ASP and GLU, the charge is computed by averaging *f* across all such residues and multiplying by their net charge of −1 at neutral pH. For basic residues such as LYS, ARG, and HIS, the charge is obtained by multiplying their protonation fraction 1 − *f* by +1 at pH = 7. We examine the charged state of the protein for various pH from our CpHMD trajectories. The overall charges of the protein(*Q*_*overall*_) are shown in Fig.1(a). At neutral pH = 7, the protein has a net positive charge of +8.36. Positive *Q*_*overall*_ increases as pH is lowered (acidic) from the neutral condition. *Q*_*overall*_ decreases with increasing pH as the medium becomes basic (pH > 7). *Q*_overall_ is positive(+1.55) at *pH* = 11.5 but becomes slightly negative (−0.44) at *pH* = 12, suggesting a charge inversion near *pH* ≈ 11.7, consistent with the previous studies.^83,84^ At extreme basic pH, *Q*_*overall*_ obtained from simulation decreases to ≈ −6.34 at pH = 13, since only the titration of acidic and basic residues has been considered in the present calculation.

**Figure 1.**
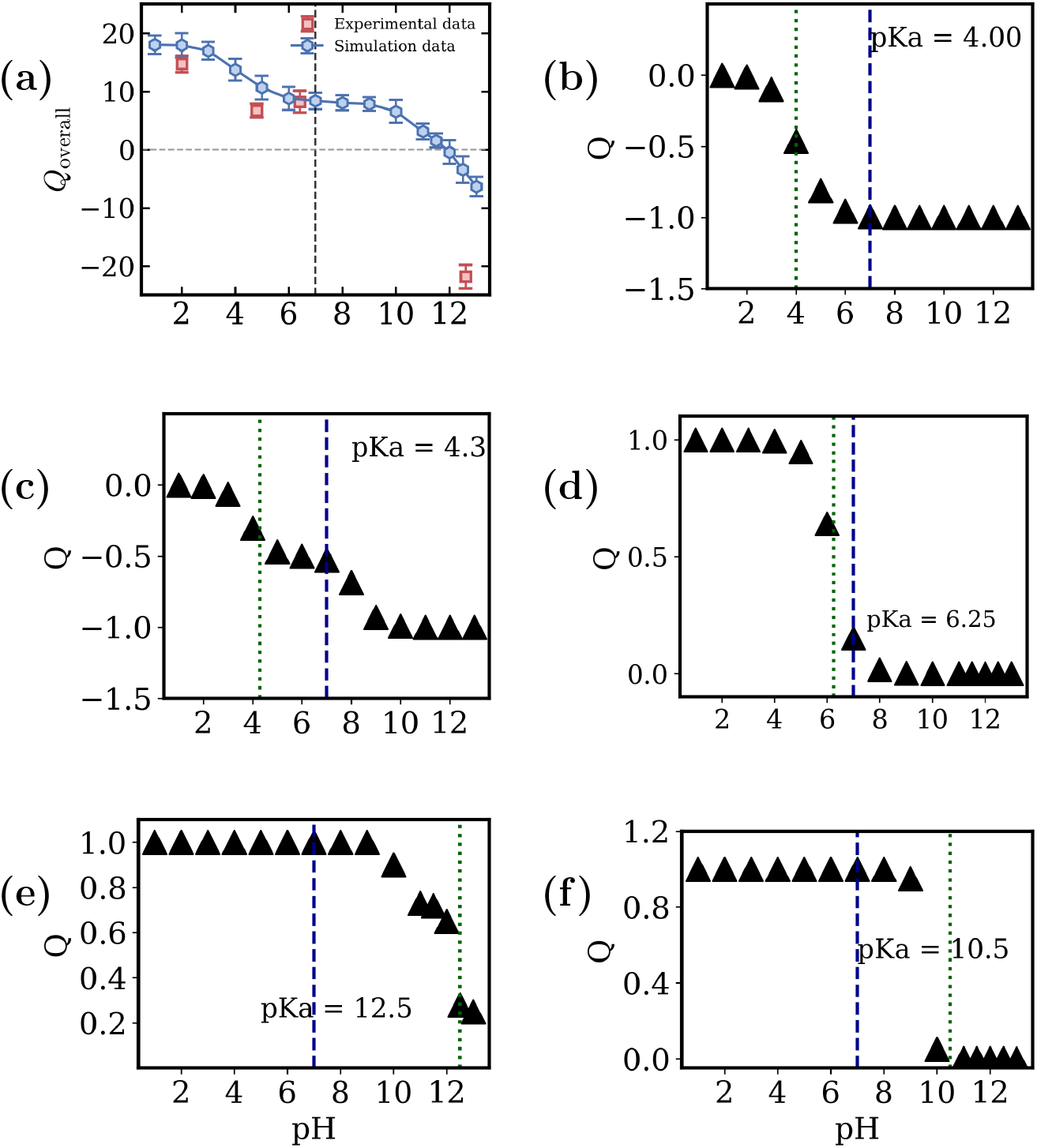
(a) *Q*_*overall*_ of lysozyme vs pH, obtained from cpHMD simulation data (blue hexagons) and experiments(red squares), (b) *Q* of ASP(7 residues). (c) *Q* of GLU(2 residues), (d) *Q* of HIS(1 residue), (e) *Q* of ARG(11 residues), (f) *Q* of LYS(6 residues) as functions of pH. Green dashed line represents the pKa values of respective residue and the blue dashed line indicates the neutral condition.

Since we are interested to capture the local changes, we consider the individual residue contribution to the overall charged state of the protein. The charge *Q* in Fig.1(b)-Fig.1(f) show the average charges of all the titrable residues. The system contains 7 ASP residues, 2 GLU residues, 1 HIS residue, 6 LYS residues, and 11 ARG residues. Fig.1(b) and Fig.1(c) show *Q* for acidic residues ASP and GLU respectively for different pH averaging over all the ASP and GLU residues. The acidic residues are negatively charged in neutral pH and remain so as pH is decreased to a more acidic medium till their pKa value is reached below which they get neutralized by loosing charge by protonation. At neutral pH, HIS is positively charged. As the pH increases beyond its pKa value, it undergoes deprotonation and becomes neutral. At extreme basic pH, HIS becomes uncharged as shown in Fig.1(d). The loss in charge by the acidic residues results in high positive *Q*_*overall*_ of the protein in this regime. The charge Q of the basic residues, averaged over all ARG and LYS respectively are shown in Fig.1(e) and Fig.1(f). They do not change their charge state in acidic condition. On the other hand, they tend to lose charge as pH increases to more basic medium beyond the neutral solvent condition. They get deprotonated beyond their respective pKa values. The charge inversion of the protein at extreme basic pH (near *pH* ≈ 11.7) arises primarily from the deprotonation of basic residues (LYS and ARG), leading to the loss of positive charge, while acidic residues (ASP and GLU) remain negatively charged.

### 3.2 Changes in protein conformation

The radius of gyration (*R*_*g*_) from the *C*_*α*_ coordinates are shown in Fig.2(a). We observe a non-monotonic behavior of protein *R*_*g*_ with changing pH. The *R*_*g*_ is minimum at neutral pH and does not change much around the neutral condition. It increases as pH is decreased in more acidic medium and increased in more basic condition. In extreme acidic and basic conditions, the protein becomes less compact than at neutral pH. This arises from changes in electrostatic interactions: in acidic conditions, protonation leads to strong electrostatic repulsion among the positively charged basic residues, promoting expansion of the protein. In contrast, under extreme basic conditions, deprotonation reduces positive charge on basic residues, weakening stabilizing interactions and making the protein less compact. It may be noted that the changes are about 6% with respect to the neutral case. This is a signature of unfolding of the protein in extreme pH conditions. Although the net charge of the protein approaches zero near the isoelectric point(pI), internal charges repel the charged residues, indicating that stability depends on charge distribution rather than net charge alone. The range of maximum stable region corresponds to the range of pH for the maximum activity of the protein.^17^

**Figure 2.**
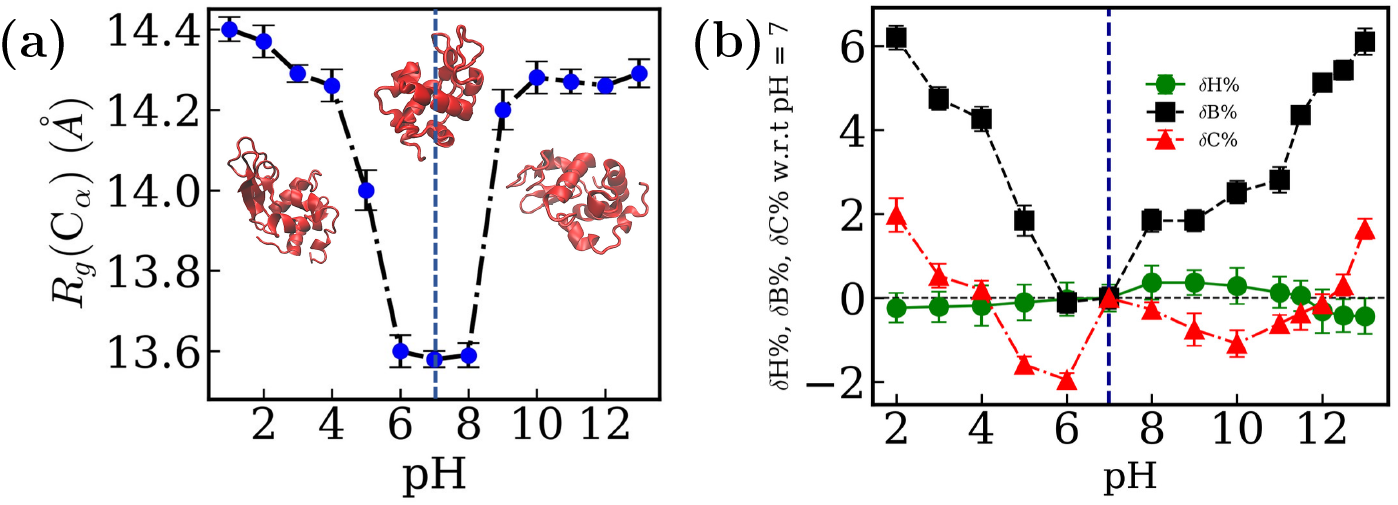
(a) *R*_*g*_ variation for different pH. (b) Percentage change in *β*-sheet(*δB*), helicity(*δH*) and random coil(*δC*) at different pH levels relative to pH = 7 (*α*-Helix: circles in green, *β*-sheet: squares in black, Coil: upper triangles in red). The blue dashed line indicates the neutral condition.

We calculate the percentages of residues having *β*-sheet, *α*-helix and random coils over equilibrated trajectories. We observe that these quantities are sensitive to pH. The percentage changes in content of beta-sheet *δB*, alpha-helix *δH* and and random coil *δC* respectively with respect to those at pH = 7 are shown in Fig.2(b). There is an increase in *β*-sheet accompanied by a decrease in the helix percentage under both acidic and basic conditions. However, the changes are small (*<* 7%). Furthermore, the Ramachandran plots for various solvent pH conditions, given in SI Fig.S3(a), Fig.S3(b), Fig.S3(c), and Fig.S3(d), support this. The small changes in the structural elements accompanied by the unfolding of the protein in extreme pH conditions, suggests the partially unfolding, consistent with the experimental results.^8–10^

### 3.3 Local changes

Our earlier studies on *α*-lactalbumin show^4^ that the structural changes occur only at localized regions in the MG state of the protein in an acidic medium. This leads us to investigate residue level changes in secondary structural elements in lysozyme in extreme acidic(pH = 3) and basic pH conditions (pH = 13). It may be noted that overall change in structural elements in these two conditions are not significant. Under extreme acidic conditions, the residues showing significant structural alterations are listed in Table.1(a) and shown over the crystal structure in Fig.3(a). Most of these are acidic residues and show large positive *δB*. A few hydrophobic and polar residues in the vicinity of the titrable acidic residues exhibit finite *δH* and *δB*. The residues undergoing changes in secondary structural elements are listed in Table.1(b), and highlighted over the crystal structure in Fig.3(b) under extreme basic conditions. We observe that several basic residues have large positive *δB*, suggesting enhancement in their tendency to form *β*-sheets. Polar and hydrophobic residues in the vicinity of these titratable sites also show noticeable secondary structure variations with significant *δB* and *δH*. Although the global structure remains largely preserved under both acidic and basic conditions, only certain residues are sensitive to pH variations. This is a signature of the molten globule(MG) state.

**Table 1.** (a) Fluctuating parts of the protein showing structural changes (in %) at pH = 3 with respect to pH = 7 and (b) pH = 13 with respect to pH = 7. (A: Acidic residues, B: Basic residues, P: Polar residues, H: Hydrophobic residues.)

| (a) |  |  |
| --- | --- | --- |
| Residue | $\delta B$ | $\delta H$ |
| GLU35(A) | 10.5 | -20.2 |
| LEU56(H) | 11.0 | 1.0 |
| ASP66(A) | 45.9 | 9.3 |
| THR69(P) | 11.8 | -23.5 |
| ASP101(A) | 21.0 | -1.8 |
| LYS116(B) | 2.2 | 16.8 |
| GLY117(P) | 3.4 | 23.0 |
| ASP119(A) | 11.0 | -8.7 |

| (b) |  |  |
| --- | --- | --- |
| Residue | $\delta B$ | $\delta H$ |
| GLU35(A) | 0.0 | -9.6 |
| SER36(P) | 0.0 | -25.0 |
| THR43(P) | -9.4 | 0.0 |
| LEU56(H) | 20.0 | 0.2 |
| ARG61(B) | 23.0 | -4.3 |
| ASP66(A) | 11.0 | 29.9 |
| GLY67(P) | 19.9 | 0.0 |
| THR69(P) | -0.2 | -23.5 |
| PRO70(H) | 26.6 | 0.0 |
| CYS80(P) | 0.0 | -18.3 |
| LEU83(H) | 23.6 | 23.2 |
| LEU84(H) | 0.0 | 24.3 |
| LYS97(B) | 32.6 | -28.0 |
| SER100(P) | 0.0 | 19.7 |
| ASP101(A) | 0.0 | 29.6 |
| GLY102(P) | 16.0 | -2.5 |
| GLY117(P) | 11.2 | 0.5 |
| ASP119(A) | 0.0 | -12.5 |
| VAL120(H) | 0.0 | 11.8 |
| ARG125(B) | 11.5 | 21.1 |
| ARG128(B) | 10.1 | 51.8 |

**Figure 3.**
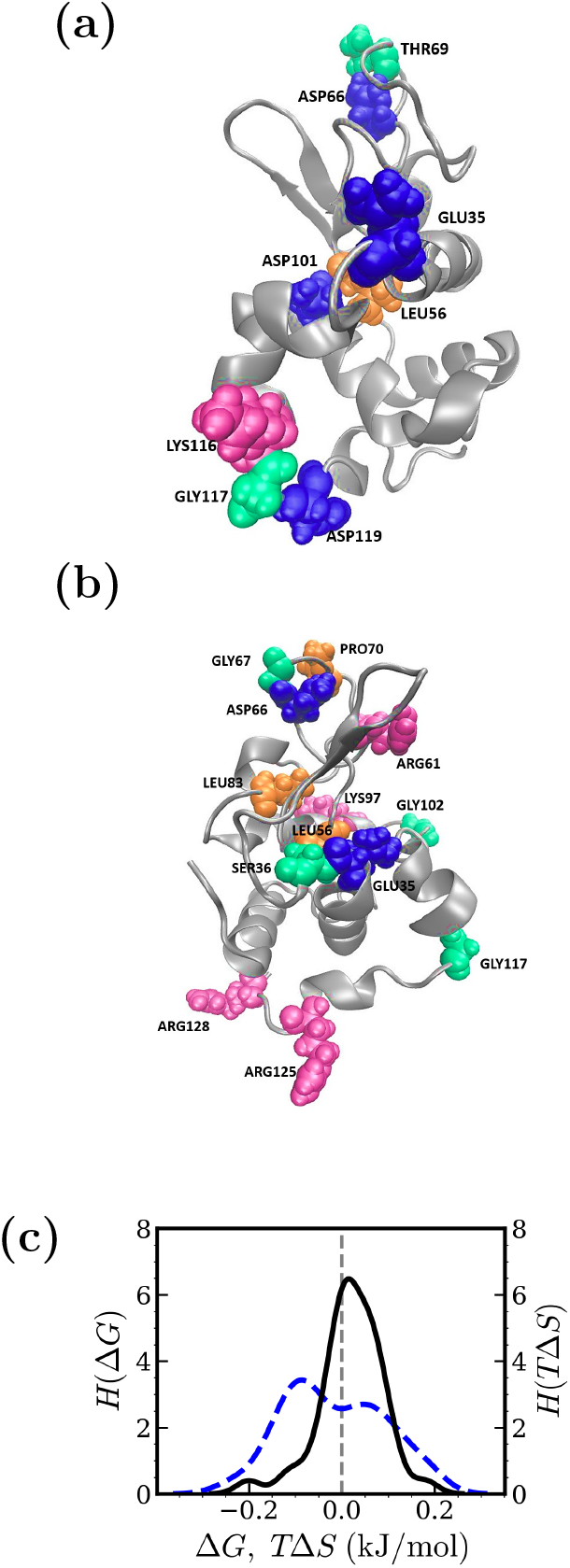
(a) Fluctuating residues at pH = 3 and (b) at pH = 13 with respect to pH = 7 are highlighted on the crystal structure,(Blue: Acidic residues, Pink: Basic residues, Green: Polar residues, Yellow: Hydrophobic residues.) (c) Histogram of Δ*G* values for all these fluctuating residues under both acidic and basic pH conditions, considering the *ϕ* and *ψ* dihedral angles, denoted in solid black line. Histogram of *T* Δ*S* values for ECs considering *ϕ* dihedral and *ψ* dihedral angles, denoted in blue dashed line.

We analyze the dihedral angle fluctuations of the protein across different pH conditions by identifying the essential coordinates (ECs) using an XGBoost-based machine learning model, followed by dPCA+ analysis combined with density based and dynamical clustering (see Methods).^4^ The top ten ECs for each pH condition are listed in Table S2(a)–S2(c) for pH = 7, 3, and 13, respectively, in the Supporting Information. Notably, the residues identified as ECs largely coincide with regions undergoing pronounced secondary structural rearrangements under extreme pH conditions, highlighting their role in pH induced conformational transitions. The corresponding free energy landscapes reconstructed from the dPCA+ analysis, together with the spatial locations of residues harboring the ECs, are shown in Fig.S4(a)–S4(f) for the different pH values. These landscapes show clear pH dependent conformational basins, with increased metastability under extreme pH conditions. This provides a direct link between changes in dihedral angle fluctuations, the contribution of key residues, and the overall conformational response of the protein to variations in pH.

We further calculate the changes in conformational free energy (Δ*G*) and entropy due to structural changes in extreme pH conditions with respect to the neutral pH as the reference structure (*T* Δ*S*)^53^ using Eq.5–Eq.7. Positive Δ*G* and *T* Δ*S* indicate residue destabilization and increased disorder, while negative Δ*G* and *T* Δ*S* values indicate greater structural stability and increased ordering relative to a reference structure. Fig.5(c) shows the distributions of Δ*G* and *T* Δ*S* over all the fluctuating residues in acidic and basic pH conditions. The peaks, centered near zero, indicate low fluctuation energy cost, consistent with previous findings.^29^ This is an indication of high conformational flexibility of these residues quantitatively.

### 3.4 Water Organization in MG state

We consider how the water organization in the vicinity of the protein surface is affected in the extreme pH conditions. The water organization around a residue is given in terms of the density profile, namely, the number of water molecules in a shell of thickness *δr* about a separation *r* between the C.O.M of the given residue to the oxygen atom of the water molecules. We investigate residue wise hydration by analyzing each titratable residue over different pH conditions. The results are shown in Fig.S5(a)–S5(z). From this analysis we observe that the residues GLU35, ASP48, ARG61, ASP66, ARG38, ASP101, LYS116, ASP119, and ARG128 exhibit noticeable changes in the surrounding water density as the pH varies. Then we average hydration surrounding the acidic, basic, hydrophobic and polar residues for different pH conditions.

*ρ*_*WA*_(*r*), namely, the average water distribution surrounding the acidic residues. *ρ*_*WB*_(*r*) denotes the water distribution surrounding the basic residues. *ρ*_*WH*_ (*r*) and *ρ*_*WP*_ (*r*) represent the average water distribution surrounding the hydrophobic and the polar residues respectively. We also compute water distribution surrounding the protein, *ρ*_*W*_ (*r*), averaged over all the residues. The hydration of the protein is quantified by the average number of water molecules within the first peak of the water density profile. Fig.4(a) shows *ρ*_*WA*_(*r*) surrounding the acidic residues with structural fluctuations. The first peak gets lower in solvent pH = 3 compared to pH = 7. This is due the protonation of these acidic residues which reduces the electrostatic interactions between the residues and the surrounding water. To the contrary the height of the first peak in *ρ*_*WA*_(*r*) is higher at pH = 13 compared to the neutral condition. This enhancement arises because the side chains of the acidic residues become deprotonated, get negatively charged and thus strengthening their interaction with surrounding water molecules. *ρ*_*WB*_(*r*) around the fluctuating basic residues is shown in Fig.4(b) for different pH conditions. We observe that the peak height of *ρ*_*WB*_(*r*) is higher in pH = 3 compared to the neutral pH condition as the basic residues carry a net positive change in extremely acidic pH. But it decreases in pH = 13 relative to pH = 7. This reduction occurs because the basic residues lose protons and become neutral, thereby weakening their interaction with water molecules. *ρ*_*WH*_ (*r*) around the hydrophobic residues (Fig.4(c)) and that around the polar residues *ρ*_*WP*_ (*r*) (Fig.4(d)) do not show any significant change in water density profile. These observations indicate that pH-driven protonation and deprotonation of residues significantly modulate electrostatic interactions, which in turn govern the hydration surrounding the protein.

**Figure 4.**
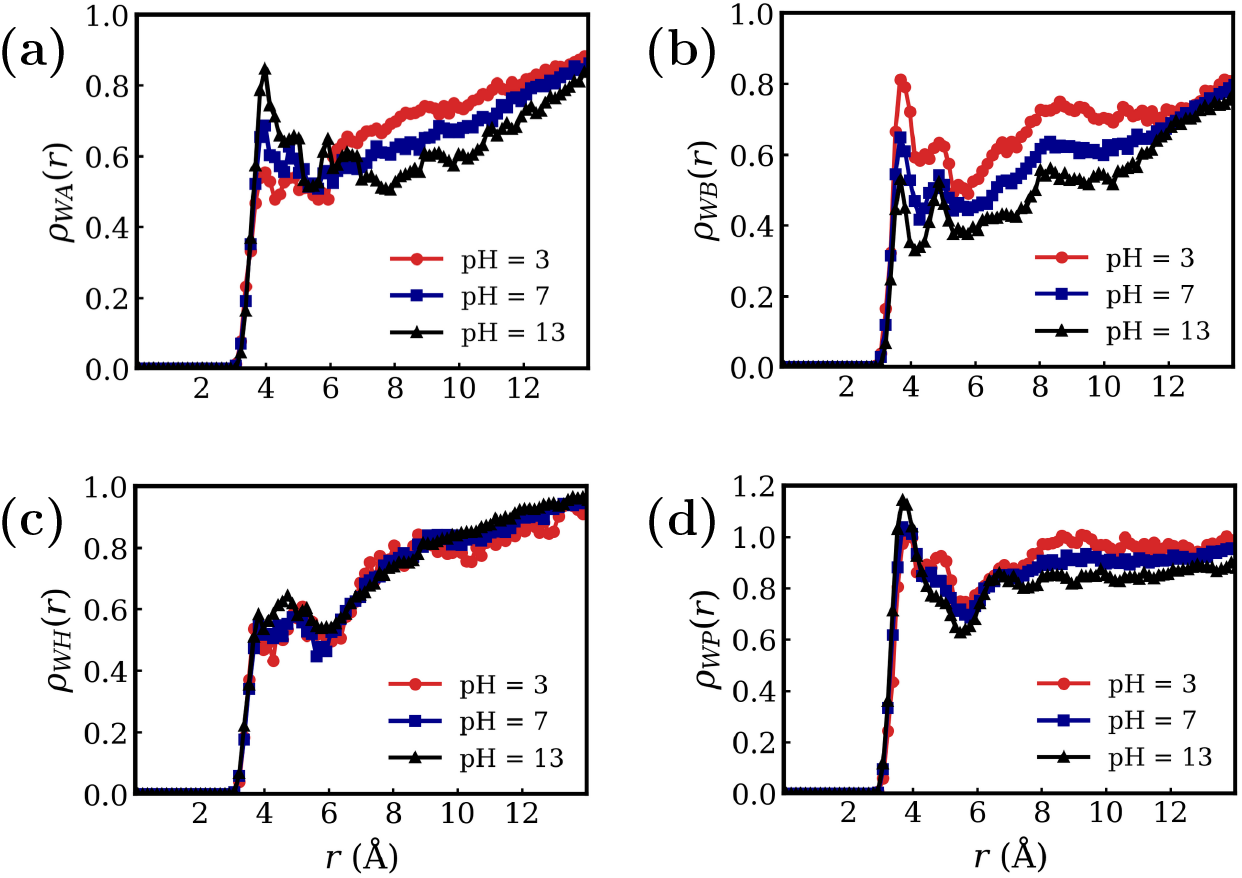
Density profile of water molecules surrounding (a) the fluctuating acidic residues(*ρ*_*WA*_(*r*)), (b) the fluctuating basic residues(*ρ*_*WB*_(*r*)), (c) the hydrophobic residues(*ρ*_*WH*_(*r*)), (d) the polar residues(*ρ*_*WP*_ (*r*)) for different pH conditions.

We also calculate the mean number of hydrogen bonds formed between the flexible residues and the surrounding hydration water molecules, *N*_*P-W*_. We observe in Fig.5(a) that *N*_*P-W*_ between acidic residues and the surrounding water molecules decreases at pH = 3 compared to pH = 7, as these residues become protonated and lose their negative charge, weakening their hydrogen bonding capability. For basic, polar, and hydrophobic residues, only slight increase in *N*_*P-W*_ is observed. At pH = 13, *N*_*P-W*_ for acidic residues increases. This is because they become deprotonated, carry negative charges while the basic residues lose their charges. This strengthens the hydrogen bonding with water molecules with the acididc residues in extreme basic pH conditions. Conversely, *N*_*P-W*_ for basic residues decreases due to deprotonation and loss of positive charge, which weakens their interaction with water in exterme pH. For polar and hydrophobic residues, *N*_*P-W*_ shows only small increases.

**Figure 5.**
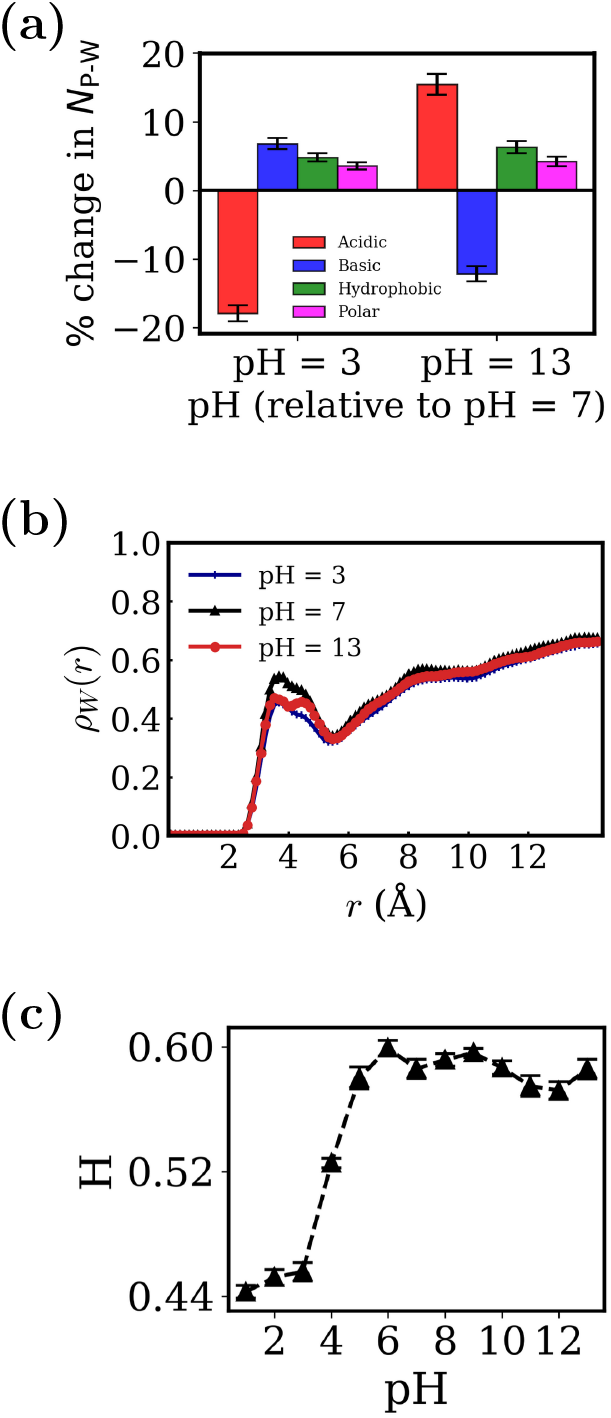
(a) Change in the number of hydrogen bonds (*N*_*P-W*_) formed between different types of residues identified as the most fluctuating regions of the protein at pH = 3 and pH = 13 with surrounding water molecules, relative to pH = 7.(Red: Acidic residues, Blue: Basic residues, Green: Hydrophobic residues, Magenta: Polar residues.) (b) The *ρ*_*W*_ (*r*) of water molecules around the protein for different pH conditions.(Vertical lines(blue): pH = 3, filled circles(red): pH = 7, upper triangles(black): pH = 13. (c) Height(H) of the first peak of *ρ*_*W*_ (*r*) for different pH conditions.

The hydration profiles of the entire protein *ρ*_*W*_ (*r*) for different pH values are shown in Fig.5(b). By numerically integrating the first peak, we obtain the overall hydration number *H*, representing the extent of water coverage around the protein surface. The variation of *H* with pH is presented in Fig.5(c). A sharp decrease in *H* is observed as the system shifts from neutral to acidic conditions, indicating reduced hydration at low pH. This arises from the protonation of the acidic residues (ASP, GLU), which lowers their negative charge and weakens electrostatic and hydrogen bonding interactions with water. In addition, increased positive charge on the protein surface under acidic conditions can promote intramolecular attractions, slightly compacting the protein and excluding water from its surface. In contrast, at higher pH values beyond pH = 7, namely, in the basic medium *H* changes in fsmall amount. Here, the deprotonation of the basic residues and the negative charge from the acidic residues, maintain a stable hydration layer around the protein.

### 3.5 Dynamics of Hydration water in MG states

We consider the dynamics of the water molecules in the hydration shell surrounding the protein in MG state. Since water molecules undergo continuous exchange between the hydration layer and the bulk phase, it is important to quantify the mean residence time *τ* of water molecules in the layer under different pH conditions. All the hydration dynamic properties are evaluated restricting to *τ*. The mean residence time, *τ*, of hydration water is determined from the decay constant of the temporal decay of the survival probability *S*(*t*) in the hydration layer. The *S*(*t*) versus *t* plots for different pH values are shown in Fig.6(a). The decay profiles are fitted with the stretched exponential form *S*(*t*) = exp[−(*t/τ*)^*β*^] where *β* is the stretching exponent. The corresponding log[− log *S*(*t*)] plots for various pH values are presented in Fig.6(b). We extract the stretching exponent *β* from the slope in Fig.6(b). *β* variation with pH is shown in Fig.6(c). The stretching exponent has a minimum around the neutral pH. The faster decay in *S*(*t*), namely, larger *β* is indicative of less electrostatic interaction between the protein surface and water molecules. The average ⟨*τ*⟩ is obtained using 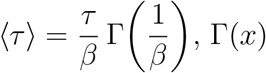 being the Gamma function. The data for different pH are given in Fig.6(d). We find that *τ* is around 40 ps at pH = 7, which is consistent with literature values.^85^ Within the range of 6 < pH < 10, *τ* remains almost constant, indicating a relatively stable hydration environment near the protein surface. However, *τ* decreases sharply under both extreme acidic and basic conditions consistent to the behaviour observed for *β*.

**Figure 6.**
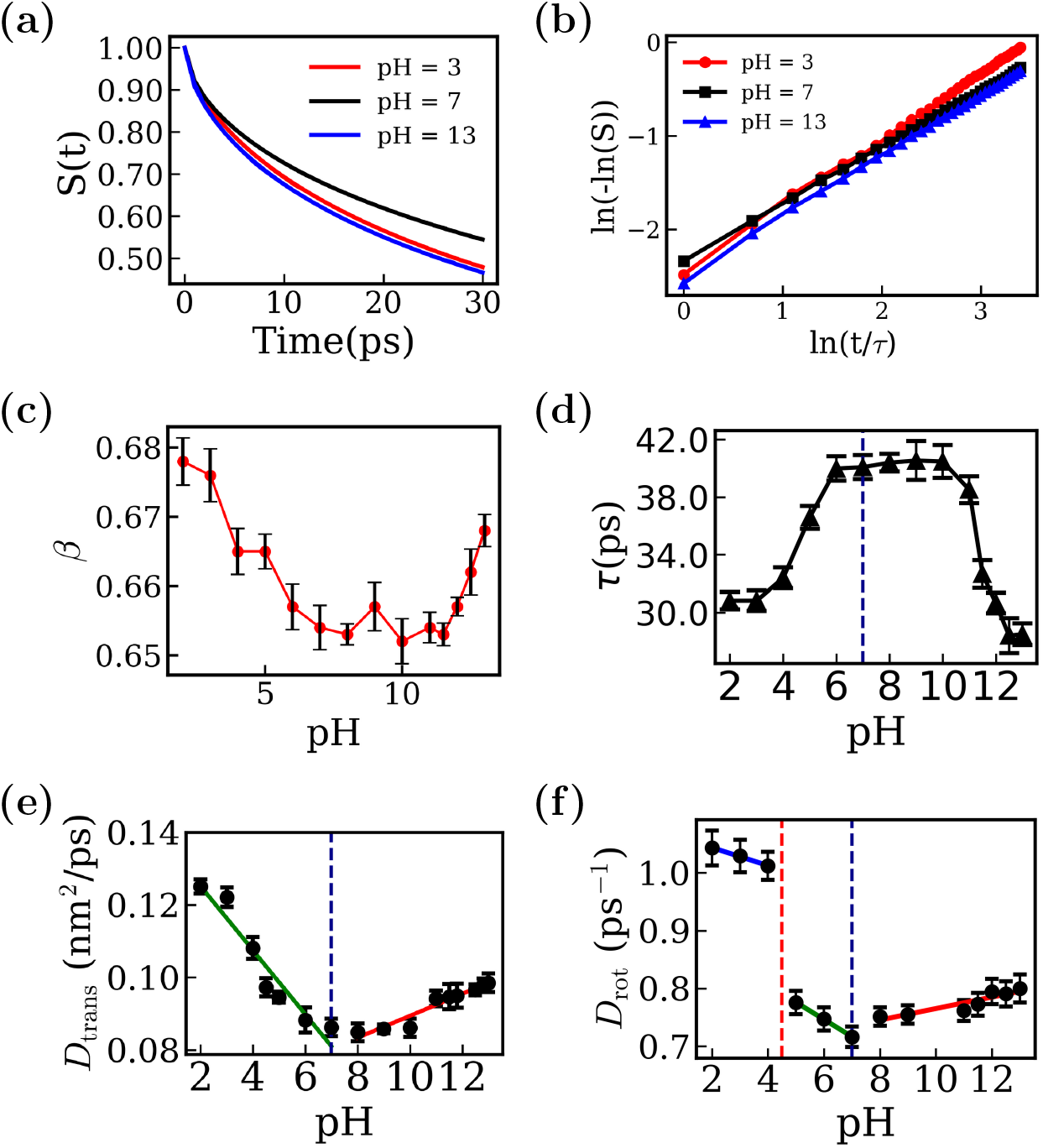
(a) Survival probability(*S*(*t*)) of hydration water for different pH conditions, (b) Plots of log[− log *S*(*t*)] over selected time intervals for each pH condition, (c) Stretched exponential parameter (*β*) from SP fitting vs. pH, (d) Residence time *τ* of hydration water for different pH conditions, (e) The *D*_*trans*_(*nm*^2^*/ps*) of hydration water for different pH conditions. The *D*_trans_ (nm^2^/ps) of hydration water shows two linear regimes: a decreasing trend for acidic to neutral pH conditions (pH ≤ 7, slope − 0.008824 (nm^2^/ps), and a weak increasing trend in the alkaline pH region (pH *>* 7, slope 0.003025 (nm^2^/ps), (f) The *D*_rot_ (ps^−1^) of hydration water dipoles exhibits three distinct linear regimes across the pH range. In Region I (2 ≤ pH ≤ 4), *D*_rot_ decreases with a slope of 0.0155 (ps^−1^). Region II (5 ≤ pH ≤ 7) shows a steeper decline with slope − 0.0298 (ps^−1^), indicating stronger slowing of rotational dynamics near neutrality. In Region III (8 ≤ pH ≤ 13), *D*_rot_ increases slightly with a positive slope of 0.0099 (ps^−1^).

We further compute the translational mean square displacement (⟨*r*^2^⟩) of the water molecules up to *τ* for a given pH averaged over initial conditions. The translational MSDs for different pH conditions are shown in Fig.S6(a). We extract the translational diffusion coefficient, *D*_trans_, from the linear regime of the MSD at long times (see Methods for details). The *D*_trans_ data are presented in Fig.6(e). We observe that the hydration water molecules diffuse faster with decrease in pH in acidic conditions. This is due to the reduced interaction between water molecules and protonated acidic residues. We observe a linear increase in *D*_trans_ with pH in the basic medium which is due to the deprotonation of basic residues.

We also calculate the rotational mean square displacement(MSD) of the hydration water dipoles within *τ* for a given pH. The rotational MSDs are shown in Fig.S6(b). We determine the rotational diffusion coefficient, *D*_rot_, from the slope of the long time linear region of the rotational MSD (see Methods). The variation of *D*_rot_ with pH is shown in Fig.6(f). The rotational diffusion is significantly faster in highly acidic environments, and increases with lowering pH till pH = 7 as observed for the translational diffusion. However, we observe two distinct linear dependencies in acidic pH region with different slopes with a break at pH ≈ 4, probably because most of the acidic residues have their pKa values near pH ≈ 4. In the basic pH region it increases slowly with pH as in the case of translational diffusion.

## 4. Discussions

We verify by experiments some of the gross simulation observations like the charge inversion and the changes in overall secondary structure. The experimental findings, illustrated in Fig.1(a), reveal the charge state of the protein across different pH values. At low pH, the protein carries a net positive charge, which persists till pH = 6.4. However, as the pH increases, a charge inversion occurs, and by pH = 12.6, the protein exhibits a net negative charge. Earlier literature reports the iso-electric point (*pI*) of the system to be approximately 10.7.^84,86^ This value is in good agreement with the results obtained from our simulations, particularly near pH ≈ 11.7 as inferred from Fig.1(a). This indicates that the computational model successfully captures the pH-dependent behavior of the system. We note that a noticeable deviation appears only in the extreme basic regime, which arises because the terminal and TYR residues are not explicitly included in the CpHMD titration scheme.

We show the frequency spectrum obtained from Mid-IR spectroscopy for pH = 6.4 in Fig.7(a). The absorbance data, using Eq.18 in the Methods section, quantifies the percentages of secondary structural elements. The peaks correspond to contributions from different secondary structure elements (e.g., *α*-helix, *β*-sheet, and random coil), and their relative changes provide insight into changes in the secondary structural elements across pH. The data, shown in Fig.7(b), indicate that under both acidic and basic conditions, beta-sheet formation increases while coil content decreases. A similar trend is observed under extreme basic pH conditions (pH = 12.6). However, these structural changes remain relatively small. Thus, despite the charge inversion the overall structure remains largely unchanged in both acidic and basic pH conditions. The experimental data support the simulation observations of charge inversion of the protein. The experimental observation on small changes in overall protein structure is also supported by the simulations.

**Figure 7.**
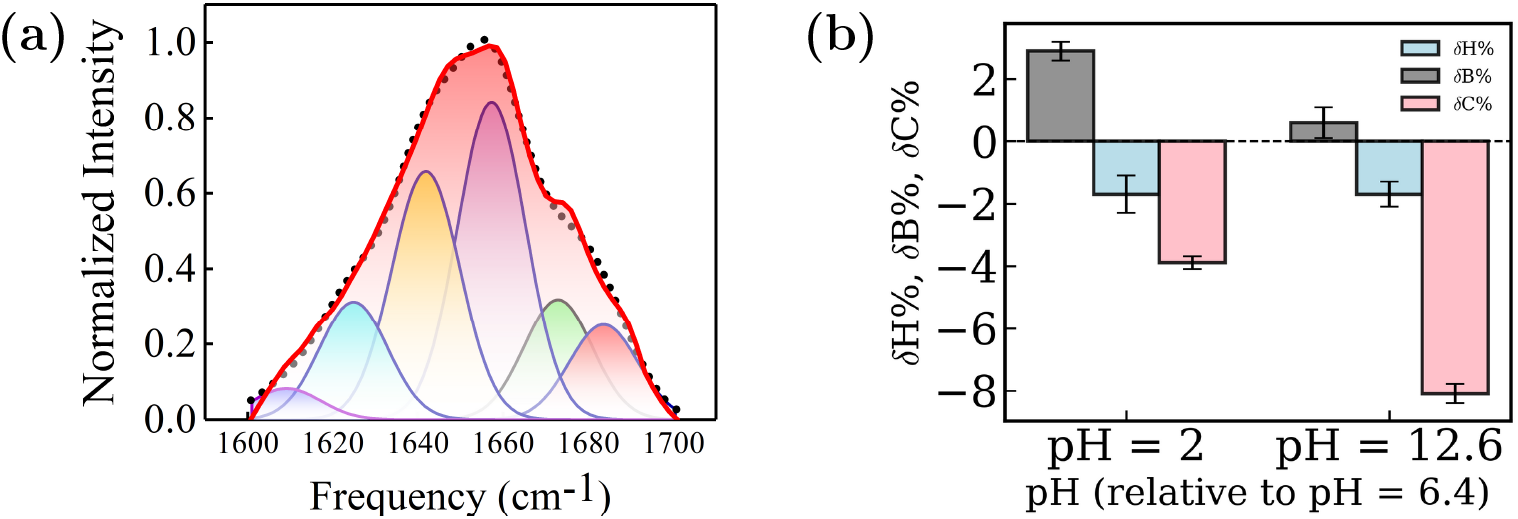
(a) Normalized intensity vs. frequency shows different components of secondary structure peaks at pH = 6.4 (Mid-IR spectroscopy)(blue: *β*-sheet(parallel), green: random coil, purple: *α*-helix, pink: *β*-strands, brown: *β*-sheet(anti-parallel)). (b) Percentage change in helicity, *β*-sheet, and random coil structures at pH = 2 and pH = 12.6 relative to pH = 6.4 (*β* sheet: bars in grey, helices: bars in blue, and coil: bars in pink).

The simulations supplement the experimental observations with more microscopic picture of the pH dependent effects: The titrable and their surrounding residues play significant role in deciding pH response of the protein and its surrounding water. Under extreme acidic conditions, acidic residues lose their negative charge due to protonation, leading to reduced electrostatic repulsion within the protein. This reduction makes these residues more solvent-exposed and capable of undergoing enhanced conformational fluctuations, suggesting an MG state. Consequently, the protein adopts a compact but more flexible structure. The fluctuating acidic residues exhibit a noticeable decrease in local water density and hydrogen bonding. In contrast, under extreme basic conditions, basic residues such as ARG and LYS lose their positive charges due to deprotonation, while acidic residues remain negatively charged. This results in an overall increase in the net negative charge of the protein and charge inversion of the protein. The protein undergoes local structural changes and partially unfolds thus indicating a MG state. The simultaneous loss of hydration from the neutralized basic residues counterbalances the enhanced hydration around negatively charged acidic residues. This leads to an overall hydration level similar to that at neutral pH.

Hydration dynamics around the protein in its MG state exhibit a clear pH dependence. The mean residence time of hydration water(*τ*), remains nearly constant between pH = 6 to 10 but decreases sharply under extreme acidic and basic conditions, reflecting faster exchange due to weakened electrostatic interactions with protonated or deprotonated residues. Both translational and rotational diffusion coefficients increase in acidic environments, indicating enhanced mobility of hydration water. In the basic regime, diffusion increases more gradually with pH. Notably, rotational diffusion shows two distinct linear regimes in the acidic range with a crossover near pH ≈ 4, consistent with the pKa values of acidic residues.

## 5. Conclusions

By integrating constant pH molecular dynamics (CpHMD) simulations, machine learning techniques, and experimental observations, we provide a microscopic analysis of pH-dependent structural and hydration changes at the molecular level in lysozyme. The primary driving force behind these pH-dependent fluctuations is the alteration of protonation states in the titrable residues with structural alterations primarily occurring at these and their adjacent residues, suggesting MG states of the protein in extreme pH conditions. The overall structure of the protein is maintained with modest changes in secondary structural elements at a local level over a wild range of pH where the overall charge of the protein changes substantially. In extreme acidic pH conditions, protonation of acidic residues leads to decreased local water density and fewer hydrogen bonds. However the basic residues remain positively charged under these conditions, promotes stronger electrostatic interactions with water and thereby exhibits comparatively enhanced hydration relative to that at neutral pH condition, although the overall change is relatively small. At basic pH, basic residues become deprotonated while acidic residues carry negative charges. In these extreme basic pH conditions, we observe a comparatively smaller change in water organization near the protein surface than at neutral pH. The translational diffusion coefficient *D*_trans_ increases linearly with decreasing pH in acidic conditions and shows a linear increase with increasing pH in the basic regime. The rotational diffusion (*D*_rot_) behaves similarly as the translational counterpart except displaying two distinct linear regimes with a crossover near pH ≈ 4. The microscopic insights reported in our studies identifying the fluctuating part of the protein along with changes in the surrounding water dynamics may be useful for drug design.

## Supporting information

Supplementary figures show RMSD/EACF, titration, Ramachandran, free energy/ECs, water density, and MSD; tables include pKa/Hill and XGBoost ECs.

## Supporting Information

Fig S1: RMSD of the system under different pH conditions; energy autocorrelation functions fitted to single-exponential decay at pH = 7, 3, and 13. Fig S2: Titration curves of titrable acidic residues showing fraction of deprotonation (*f*) as a function of pH. Fig S3: Ramachandran plots of lysozyme at pH = 3, 7, 11, and 13. Fig S4: Free energy landscapes from dPCA+ analysis and the corresponding essential coordinate (EC) residue mapping on the crystal structure at pH = 7, 3, and 13. Fig S5: Density profiles of water surrounding each titrable residue at different pH conditions. Fig S6: Translational mean square displacement (⟨Δ*r*^2^(*t*)⟩) and rotational mean square displacement (⟨Δ*ϕ*^2^(*t*)⟩) of hydration water at different pH conditions. Table S1: Comparison of computed p*K*_*a*_ values with experimental values and corresponding Hill coefficients for all titrable residues. Table S2: Top ten essential coordinates (ECs) identified using XGBoost at pH = 7, 3, and 13.

## 6. Acknowledgements

We sincerely acknowledge Dr. Indrani Bhattacharya for her support in our experiments. We also thank Dr. Abhik Ghosh Moulick, Dr. Anirban Paul, Dr. Kanika Kole, Mr. Pritam Roy, Mr. Bidhan Kumbhakar and Mr. Sabuj Mandal for their valuable discussions. AS thanks DST(Department of Science and Technology, GOI) for funding through the Ph.D. program and SNBNCBS, Kolkata for providing the computational facility. JC thanks the CSIR for financial support through the EMR scientists scheme. We thank DST(Department of Science and Technology, GOI) for funding through the Ph.D. program and SNBNCBS, Kolkata for providing the computational facility. We also acknowledge National Supercomputing Mission (NSM) for providing computing resources of ‘PARAM RUDRA’ at S.N. Bose National Centre for Basic Sciences, which is implemented by C-DAC and supported by the Ministry of Electronics and Information Technology (MeitY) and Department of Science and Technology (DST), Government of India.

---

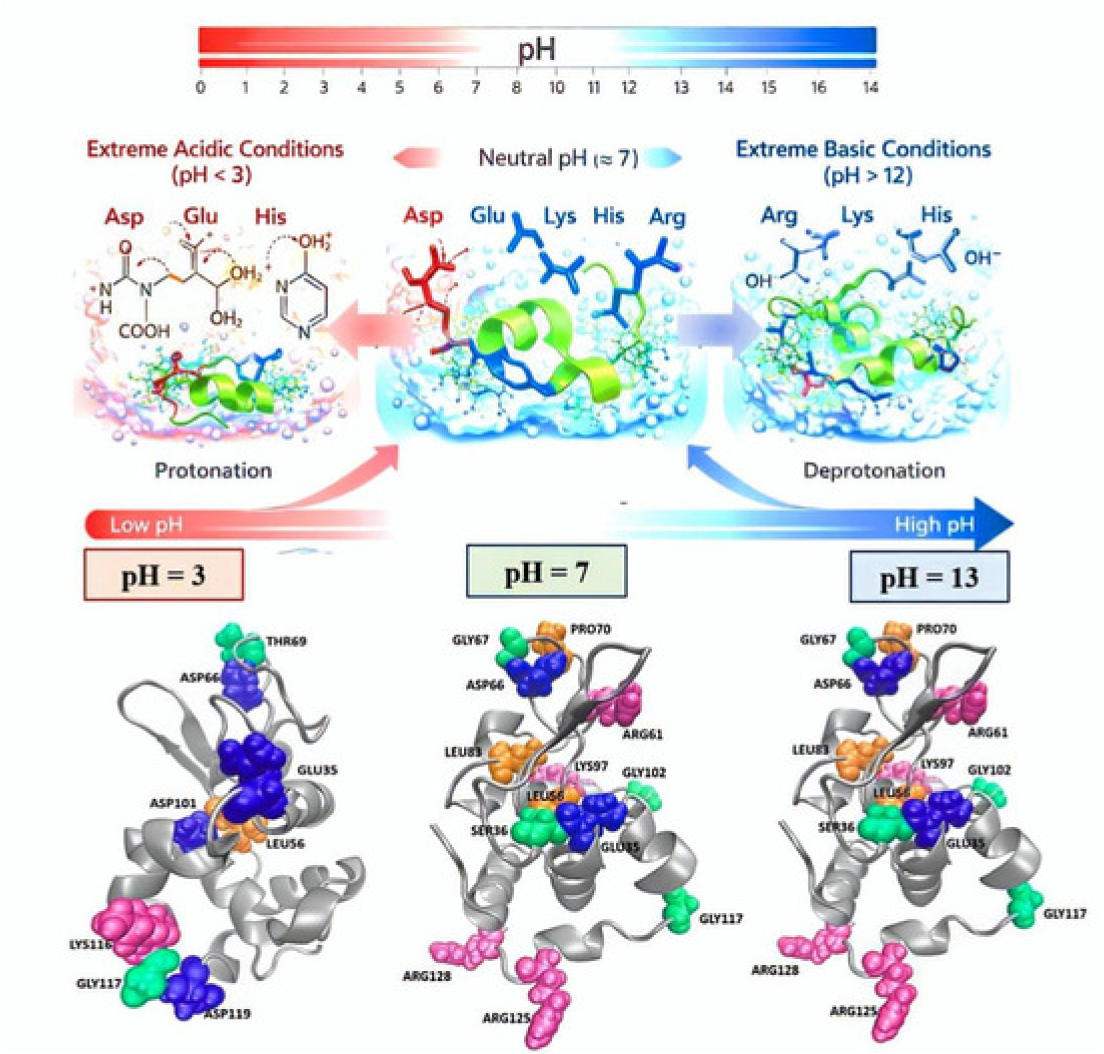

