## Supplementary figures show RMSD/EACF, titration, Ramachandran, free energy/ECs, water density, and MSD; tables include pKa/Hill and XGBoost ECs. for "pH Induced Changes in Protein Structure and Hydration"


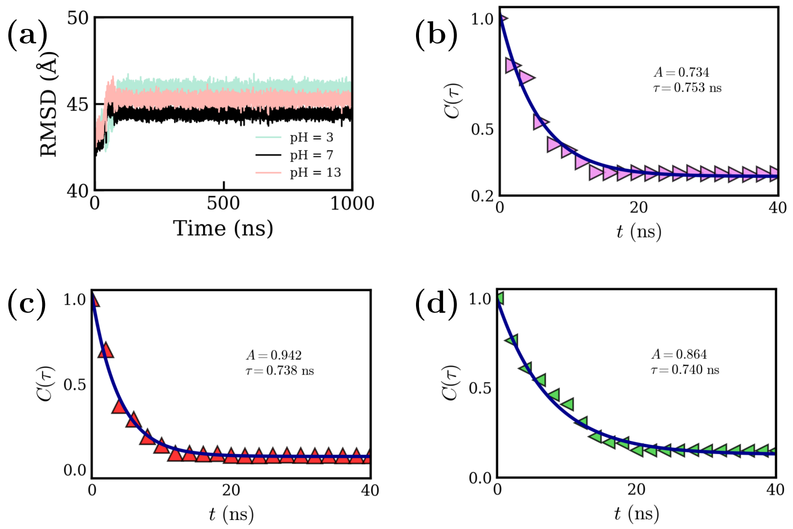


**Figure S1:** (a) RMSD of the system for different pH conditions , Energy autocorrelation functions calculated from the equilibrated trajectories and fitted to a single-exponential decay function for (b) pH = 7, (c) pH = 3, (d) pH = 13.

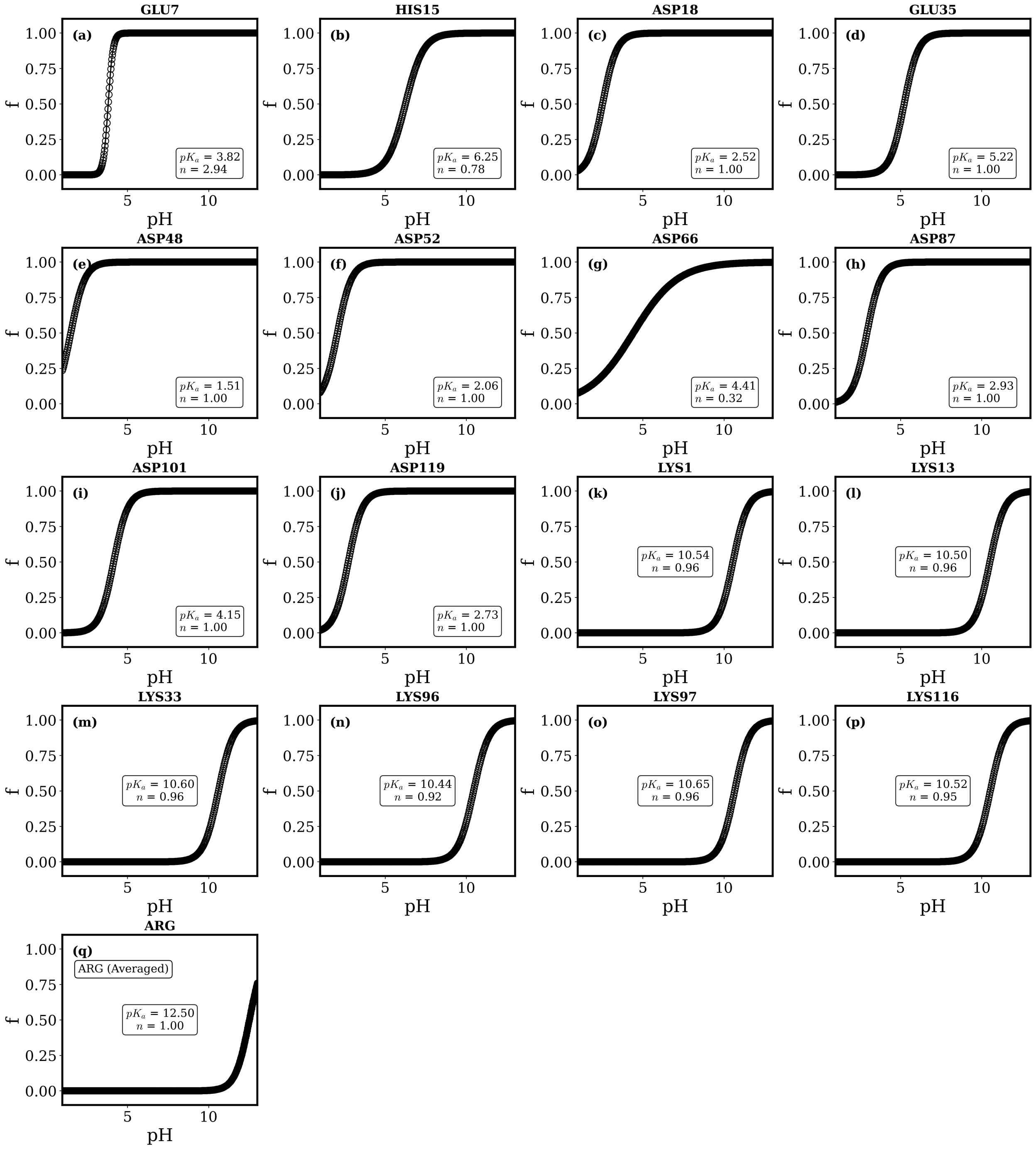


**Figure S2:** (a) - (q) The Titration curve corresponding to each titrable acidic residues showing fraction of de-protonation($f$) over a range of pH.

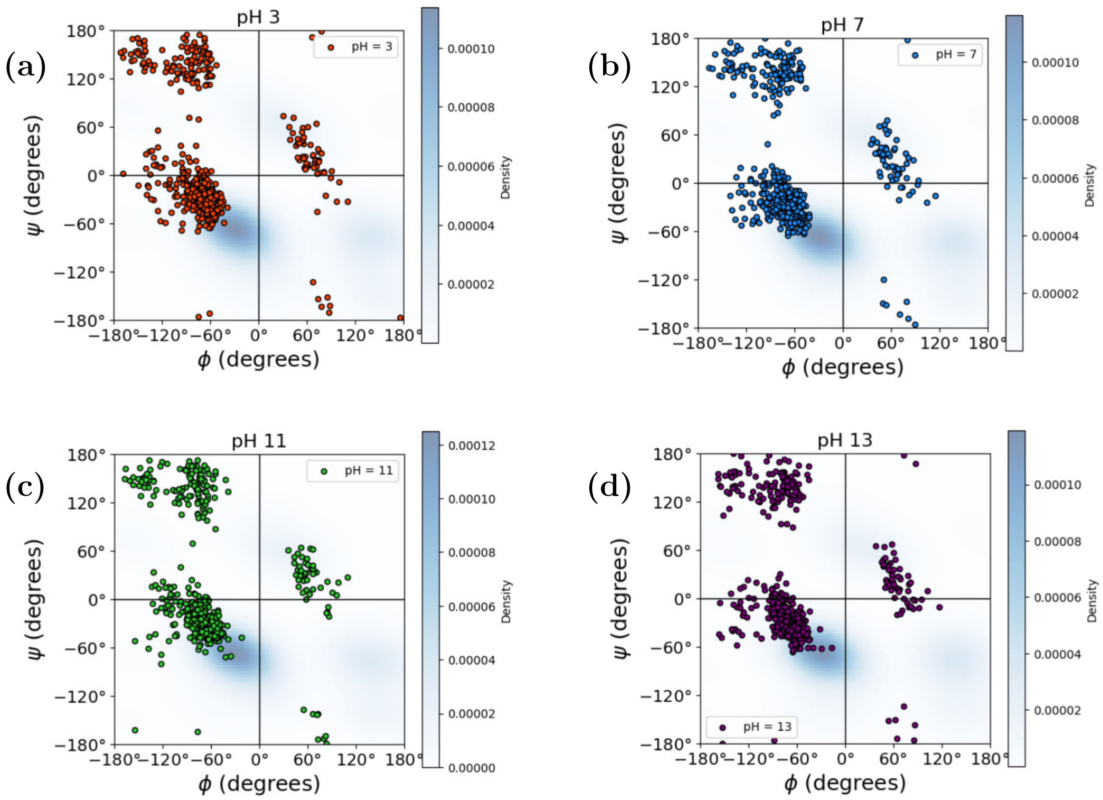


**Figure S3:** Ramachandran plots of Lysozyme at different pH values: (a) pH = 3, (b) pH = 7, (c) pH = 11, and (d) pH = 13.

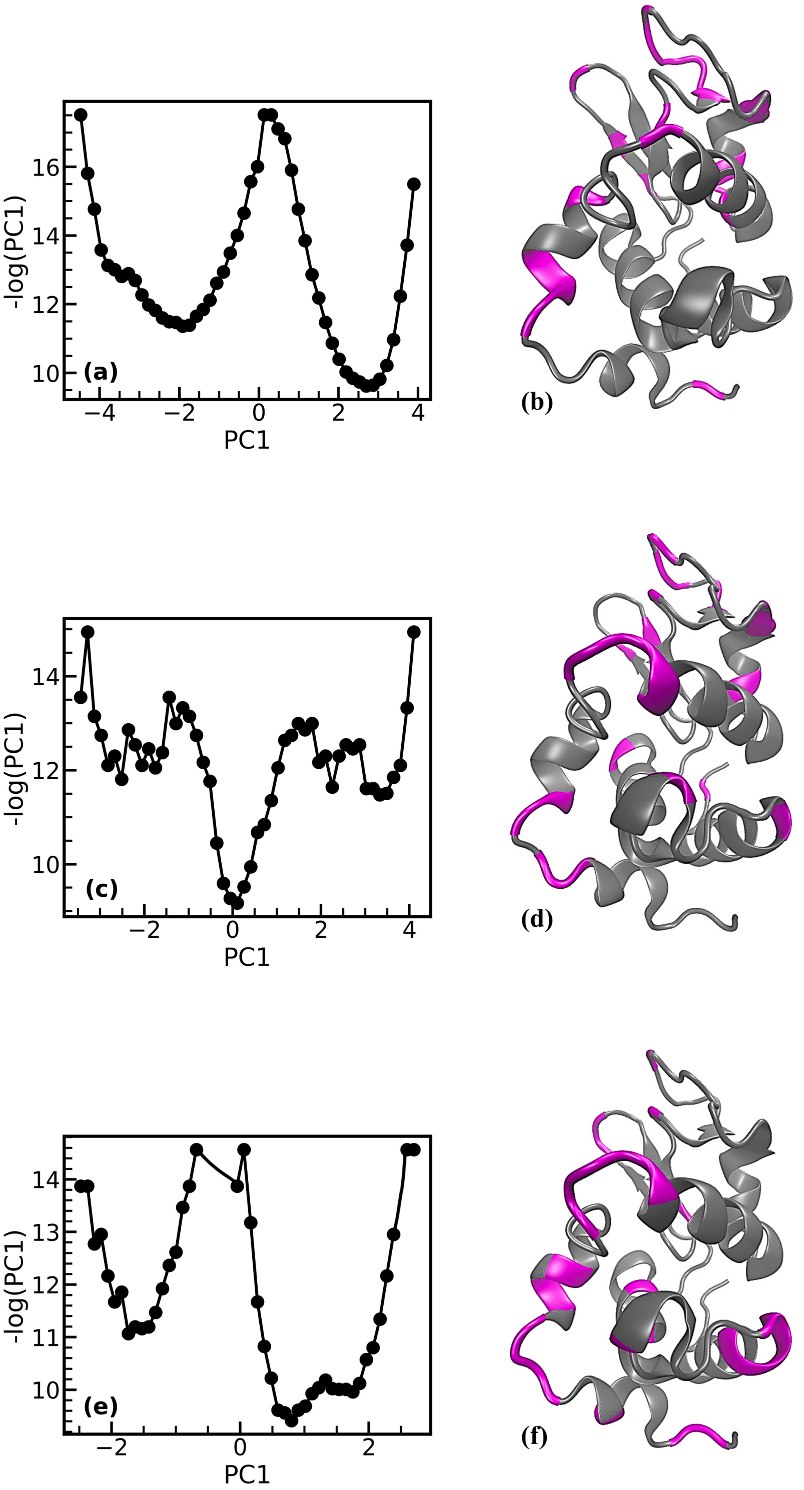


**Figure S4:** (a) Free energy landscape obtained from dPCA+ for pH = 7 showing metastability, (b) The residues harbouring the ECs highlighted in deep magenta on the crystal structure for solvent pH = 7, (c) Free energy landscape obtained from dPCA+ for pH = 3 showing metastability, (d) The residues harbouring the ECs highlighted in deep magenta on the crystal structure for solvent pH = 3, (e) Free energy landscape obtained from dPCA+ for pH = 13 showing metastability, (f) The residues harbouring the ECs highlighted in deep magenta on the crystal structure for solvent pH = 13.

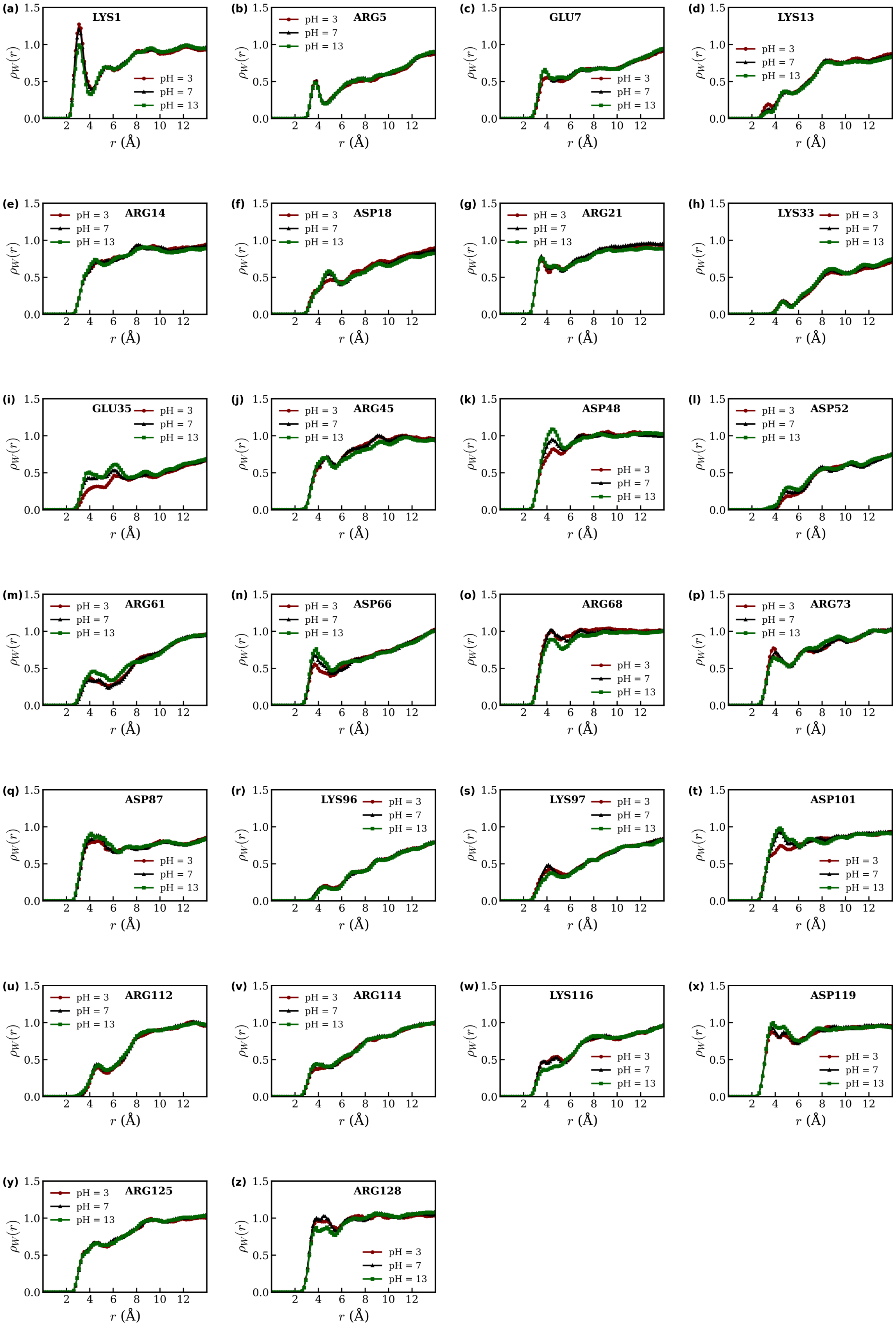


**Figure S5:** Density profile of water surrounding each titrable residue for three different pH conditions

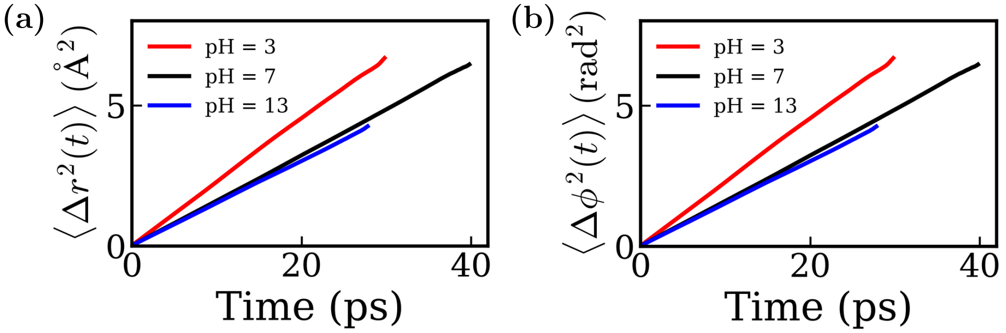


**Figure S6** (a) Translational MSD($\langle \Delta r^2(t) \rangle$) of hydration water for different pH conditions, (b) Rotational MSD ($\langle \Delta \phi^2(t) \rangle$) of hydration water for different pH conditions.

| Residue | Computed pKa | Experimental pKa | Hill coefficient |
|---|---|---|---|
| GLU7 | 3.82 | 2.6 | 2.94 |
| HIS15 | 6.25 | 5.5 | 0.78 |
| ASP18 | 2.52 | 2.8 | 0.52 |
| GLU35 | 5.22 | 6.1 | 0.18 |
| ASP48 | 1.51 | 1.4 | 0.1 |
| ASP52 | 2.06 | 3.6 | 0.32 |
| ASP66 | 4.41 | 1.2 | 0.32 |
| ASP87 | 2.93 | 2.2 | 0.52 |
| ASP101 | 4.15 | 4.5 | 0.76 |
| ASP119 | 2.73 | 3.5 | 0.43 |
| LYS1 | 10.54 | 10.8 | 0.96 |
| LYS13 | 10.5 | 10.5 | 0.96 |
| LYS33 | 10.6 | 10.5 | 0.96 |
| LYS96 | 10.44 | 10.8 | 0.92 |
| LYS97 | 10.65 | 10.5 | 0.96 |
| LYS116 | 10.52 | 10.2 | 0.95 |
| ARG5 | 12.5 | 13.8 | 1.0 |
| ARG14 | 12.5 | 13.8 | 1.0 |
| ARG21 | 12.5 | 13.8 | 1.0 |
| ARG45 | 12.5 | 13.8 | 1.0 |
| ARG61 | 12.5 | 13.8 | 1.0 |
| ARG68 | 12.5 | 13.8 | 1.0 |
| ARG73 | 12.5 | 13.8 | 1.0 |
| ARG112 | 12.5 | 13.8 | 1.0 |
| ARG114 | 12.5 | 13.8 | 1.0 |
| ARG125 | 12.5 | 13.8 | 1.0 |
| ARG128 | 12.5 | 13.8 | 1.0 |

**Table S1:** Comparison of the computed pka values with the experimental pka and the corresponding fitted Hill coefficient for all titrable residues.

(a)

| Residue id | Dihedral Angle | Residue Name |
|---|---|---|
| 116 | Ψ | GLY |
| 68 | Φ | ARG |
| 60 | Ψ | SER |
| 48 | Φ | ASP |
| 117 | Ψ | GLY |
| 101 | Φ | ASP |
| 87 | Φ | ASP |
| 117 | Φ | GLY |
| 115 | Φ | CYS |
| 66 | Ψ | ASP |

(b)

| Residue id | Dihedral Angle | Residue Name |
|---|---|---|
| 35 | Ψ | GLU |
| 66 | Ψ | ASP |
| 101 | Φ | ASP |
| 119 | Ψ | ASP |
| 69 | Φ | THR |
| 116 | Ψ | LYS |
| 102 | Ψ | GLY |
| 102 | Φ | GLY |
| 101 | Ψ | ASP |
| 117 | Φ | GLY |

(c)

| Residue id | Dihedral Angle | Residue Name |
|---|---|---|
| 102 | Ψ | GLY |
| 70 | Ψ | PRO |
| 128 | Ψ | ARG |
| 66 | Ψ | ASP |
| 97 | Φ | LYS |
| 67 | Φ | GLY |
| 83 | Φ | LEU |
| 61 | Φ | ARG |
| 115 | Ψ | CYS |
| 125 | Ψ | ARG |

**Table S2:** (a) First ten ECs obtained from XGBoost analysis for pH = 7, (b) First ten ECs obtained from XGBoost analysis for pH = 3, and (c) First ten ECs obtained from XGBoost analysis for pH = 13.
